# Mutational landscapes of normal breast during age and pregnancy determine cancer risk

**DOI:** 10.1101/2020.09.04.277715

**Authors:** Biancastella Cereser, Neha Tabassum, Lisa Del Bel Belluz, Angela Yiu, Sladjana Zagorac, Cristian Miere, Benjamin Werner, Nina Moderau, Alicia Rose Jeffries-Jones, Justin Stebbing

## Abstract

The accumulation of somatic mutations in the healthy breast throughout life and pregnancy is poorly understood^1–10^. Similarly, the mutational landscape of both epithelial and stromal components of the mammary gland has not been investigated. Both are relevant for breast cancer (BC), as the interplay between age, pregnancy, and cancer risk has not been fully characterized^11^. We describe whole genome sequencing analysis of epithelial and stromal compartments from the normal breast. We show that, in a similar way to other normal organs, the mutational burden of the mammary nulliparous epithelium significantly increases with age. In a nulliparous status, mutated clones are maintained at a consistently small size throughout the life of the individual; however, at parity, pre-existent clones significantly increase in size with age. Both epithelial and stromal compartments of the healthy breast contain pre-existing known cancer mutations, albeit at low rate, indicative of subsequent positive selection for mutations in tissue-specific driver genes. In line with this, both compartments also present gene enrichment in preferentially mutated cancer pathways. Our results show that mutational landscapes differ between the parous and nulliparous epithelium and suggest an explanation for both differential breast cancer risk and development of pregnancy-associated BC (PABC).

## INTRODUCTION

In all adult healthy tissues, somatic mutations increase through adult life^1–10^, but the mutational landscape of the human breast has not been characterized thus far. Reasons for the lack of in-depth sequencing data include the variation of the mutational burden observed amongst sampled individuals due to the different clonal expansion of the mammary epithelium during pregnancy and difficulties of working with breast tissue considering the prevalence of the adipocyte component.

Compared to other solid tissues, the adult human nulliparous breast is characterized by a generally slow proliferating epithelium. A recent study estimates the gain of 0.7 ± 0.1 new cells per day in pre-menopausal women^12^, and the proliferation index can increase nearly 5 times during pregnancy^13^. Furthermore, after breast-feeding, a strong apoptotic drive, which determines the beginning of post-partum mammary involution^14^, contributes to major tissue remodeling which can last up to 10 years after parturition^15^. Several studies have recognized the complex contribution of parity and age to the mammary landscape, in particular to the risk of BC development^11,16,17^. Women who reach full-term pregnancy before the age of 24 have a lower long-term BC risk compared to age-matched nulliparous women^11,16^. On the other hand, women who give birth for the first time after age 35 are more likely to develop BC than early-pregnancy age-matched groups^16^. Furthermore, for both age groups, shortterm breast cancer risk significantly increases after each pregnancy, seen in the subset of PABC^18–21^.

While previous research characterized the genetic composition of BC detected during pregnancy^22^ or of BC samples collected from women with previous pregnancies of different ages^22^, here we sequence for the first time the whole genome of the healthy breast samples from 25 individuals and determine how both age and pregnancy alter the genetic composition of the mammary gland, both in the epithelial and stromal compartments.

## RESULTS

### Sample information and sequencing

We analyzed a total of 25 frozen healthy breast tissues. Patients with no previous use of hormonal contraception and with uniparity were preferentially included in the study (n uniparous = 15/17). See **Supplementary Table 1** for clinical information. To increase the detection of mutated cells present at low frequency in the samples, we laser-captured both stroma and epithelium individually, to avoid cross-contamination of the two cellular compartments (**Fig. 1a-c**). We also analyzed further 4 parous individuals as an additional subset, as shown in the discussion section.

**Figure 1.**
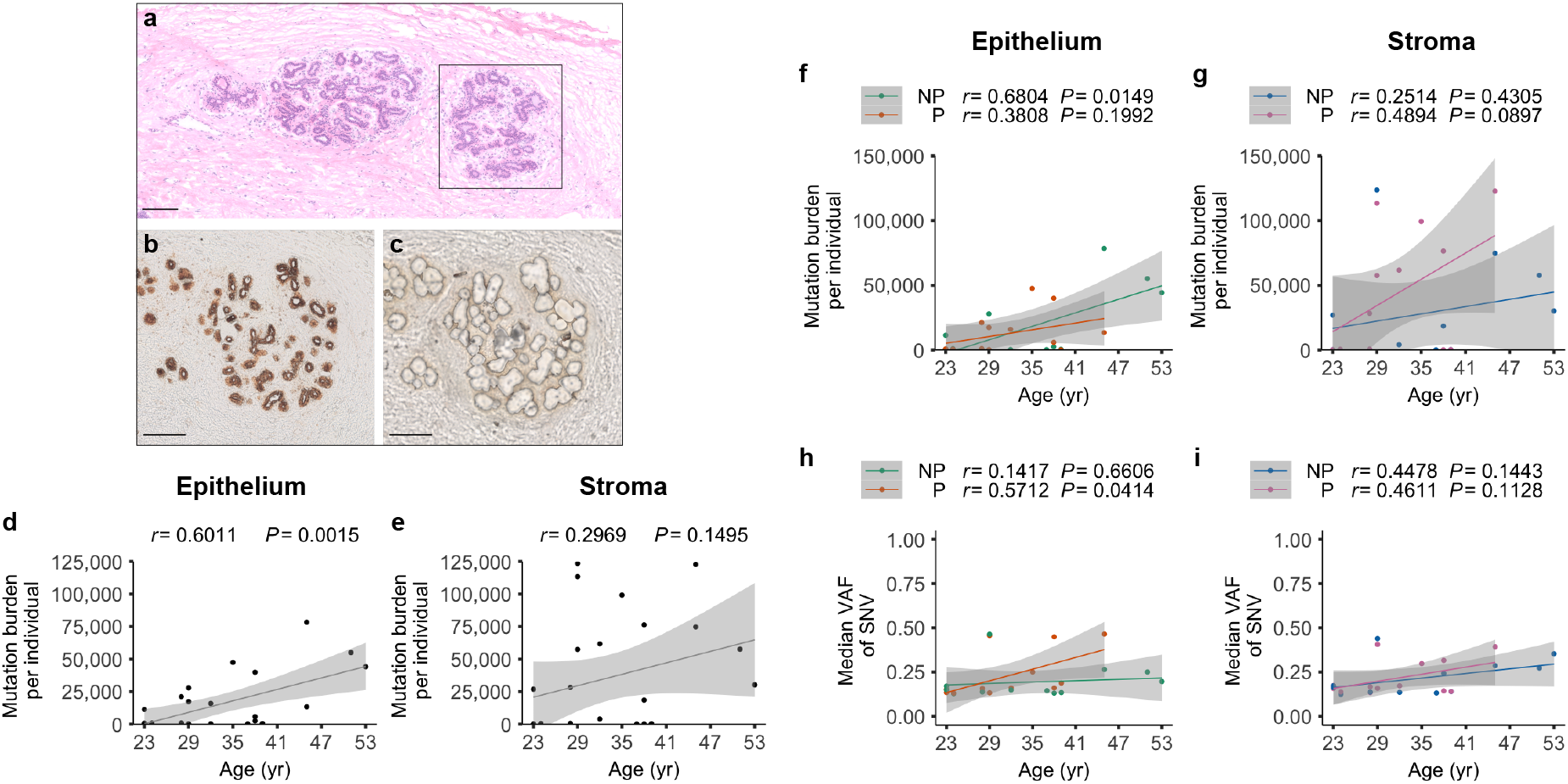
Correlation of the mutation burden with age of collection and parity in the healthy breast. **(a-c)** Representative images of non-continuative serial sections of a healthy mammary gland from a 29-year-old nulliparous woman. **(a)** Hematoxylin and eosin staining showing two terminal ductular lobular units (TDLU) surrounded by stroma. **(b)** Pre- and **(c)** post-laser-capture microdissection of the TDLU highlighted in **(a)** The section was first stained with an enzymatic reaction, which reacts to the activity of the mitochondrial cytochrome C oxidase and allow easy identification of the epithelial cells. Scale bar = 150μm. **(d,e)** Mutation burden shown as one dot for each donor (n = 25 donors, some dots overlapping). Pearson correlation (*r*) with age and P values (*P*) from linear regression are shown for the **(d)** epithelial and **(e)** stromal compartment. **(f,g)** Mutation burden and **(h,i)** median variant allele frequency of SNV shown as one dot for each donor (n = 13 parous, n = 12 nulliparous donors, some dots overlapping) in the **(f,h)** epithelial and **(g,i)** stromal compartment. Pearson correlation (*r*) with age and P values (*P*) from linear regression are shown for each group (NP = nulliparous, P = parous).

The main challenge of using normal breast for this type of study is the paucity of epithelial cells, therefore we resorted to the analysis of one sample per donor. The median sequencing depth was 40.9X and all samples had a depth of >15X, with 90% of the samples characterized by a depth of >30X (**Supplementary Table 2**).

### Variation of the mutational burden of the mammary gland with age

Overall, in both epithelium and stroma, the somatic mutational burden increased with age of collection (**Fig. 1d,e**), with an estimated rate of approximately 1470 mutations/year (muts/year). A significant accumulation with age was seen uniquely in the epithelium (*P* = 1.5×10^-3^) and could be due to a lower inter-patient variability of the number of mutations accumulated compared to the stromal compartment.

In the epithelia, the mutational burden in the 25 individuals varied amongst samples, ranging between 75 and 32,266 substitutions (median 1,866) and between 20 and 49,365 indels (median 816). In the stroma, we also observed a range between 74 and 35,976 substitutions (median 9,425) and between and 5 and 91,767 indels (median 18,739) (**Fig. 1d,e**, **Extended Data Fig. 1a-d** and **Supplementary Table 3**). There was no correlation between burden and sequencing coverage amongst the samples.

Significant positive correlations with age were identified in the epithelium for the individual mutational burden of synonymous and non-synonymous mutations. However, in the stromal compartment, only non-synonymous mutations significantly accumulated with age in the healthy breast tissue (**Extended Data Fig. 1e-h**).

### Mutational burden and clonality of the healthy breast in relation with age and pregnancy

Epidemiologic studies have determined a double role of pregnancy in the contribution of BC risk. Our aim was to detect how not only age but also pregnancy may affect the mutational burden in the healthy breast. We define the parity status as the phase of tissue remodeling, which starts from conception and has an effect up to 10 years after pregnancy^15^.

In the epithelium, the mutational burden differed according to parity. Only in the nulliparous breast the burden significantly increased with age compared to the parous counterpart (*P* = 1.5×10^-3^). The rate of accumulation was approximately 1740 muts/year in the nulliparous gland, an 8-fold decrease compared to a parous gland at 25 years of age, but a 1.55-fold increase compared to a parous gland at 50 years of age (**Fig. 1f** and **Extended Data Fig. 1i-l**). Remarkably, compared to the epithelium, the stromal compartment accumulates mutations in a different mode. While not significant, parous individuals tend to accumulate more indels and synonymous mutations (defined as in Methods section) with age compared to the nulliparous samples (**Fig. 1g** and **Extended Data Fig. 1m-p**). We estimate the rate of accumulation of mutations is approximately 934 muts/year in the nulliparous breast, similar to the parous gland at 25 years of age, but a 1.55-fold increase from a parous gland at 50 years of age (**Fig. 1g**). A specific age at which parity influences this trend is not evident from the stromal analysis, suggesting the contribution of pregnancy to BC risk is attributable mainly to changes in the epithelium.

Similarly, to the number of mutations, there was high variance of the respective variant allele frequencies amongst individuals in both cellular compartments (**Fig. 1h,i** and **Extended Data Fig. 2**). In the epithelium, the median total clone size tends to increase with age irrespective of parity, but this is particularly evident for the parous group (*P* = 4.1×10^-2^), compared to the nulliparous group, which did not show the same increase (*P* = 6.6×10^-1^) (**Fig. 1h**). We did not observe the same trend in the stromal compartment of the parous individuals.

These data indicate that later parity (more than 30 years of age) results in a lower mutational burden of the mammary epithelia, but higher clonal expansion compared to a status of nulliparity of the same age.

### Mutational burden within breast-cancer associated genes

As age of pregnancy is a risk factor for both PABC and post-menopausal BC, we analyzed the mutational burden and clonality of 93 BC-associated genes, as previously identified from 560 breast cancer whole genome sequences^23^.

We detected somatic mutations in all 93 driver genes in all healthy patients, albeit with different frequency, in both the epithelium and stroma. Some of these events were identical and occurring in both compartments. This is in line with the findings from other healthy tissues, which all confirm the presence of mutations in cancer-associated genes in the normal epithelium and blood ^1,3–7,24^. However, in the breast, there is preferential accumulation of mutations in *ESR1* and *RUNX1*, mutated in approximately 60% of samples in the parous and nulliparous group, respectively (**Fig. 2a,b and Supplementary Table 4**). We detected non-synonymous mutations in *ERBB2* and *PTEN* in 12% of the epithelial samples (3/25), and in *NOTCH2* and *PTEN* in the same percentage of distinct stromal samples (**Extended Data Fig. 3a,b**).

**Figure 2.**
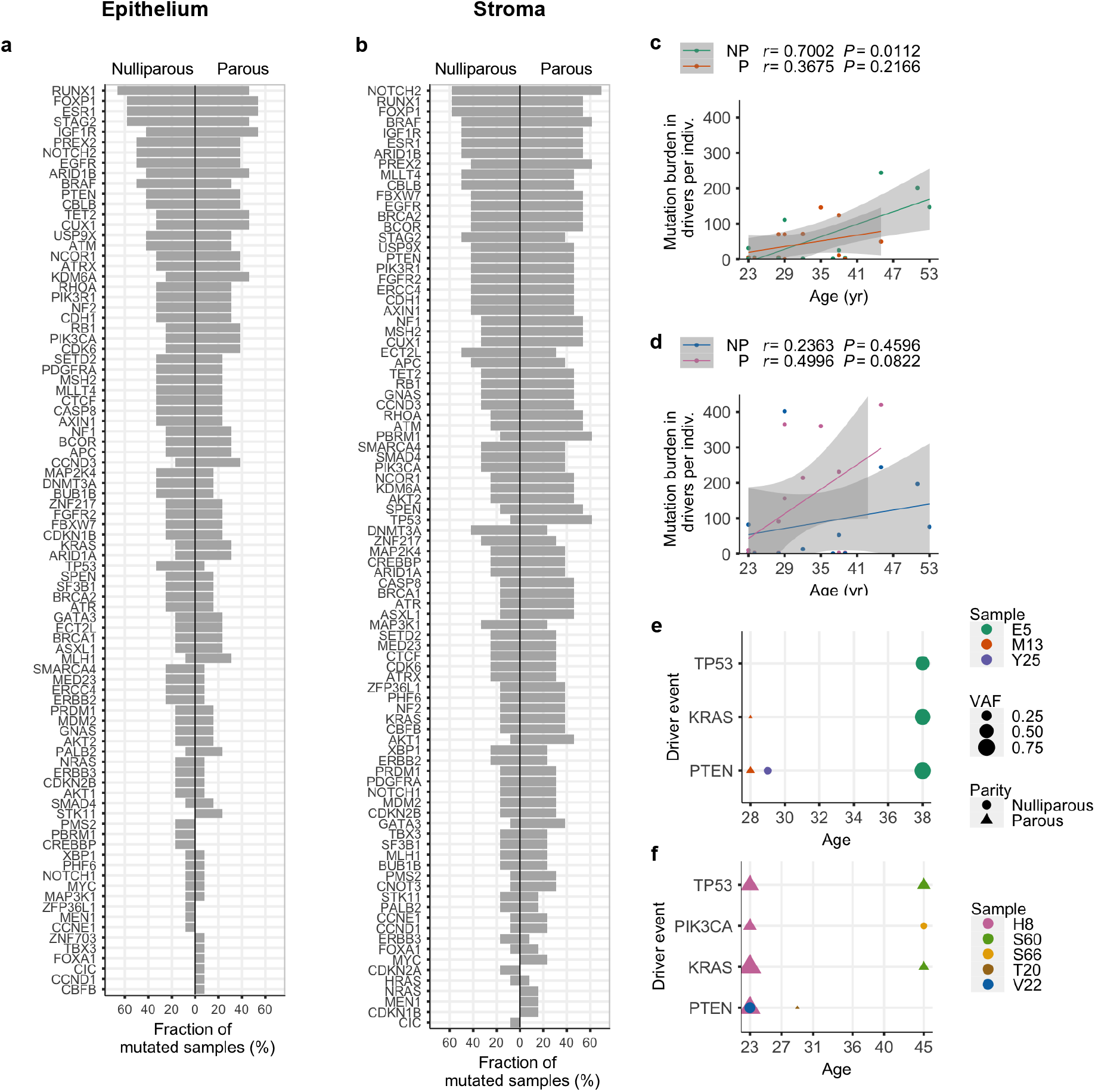
Mutation burden of driver genes in the healthy parous and nulliparous breast. **(a,b)** Fraction of nulliparous (n = 12) and parous (n = 13) samples carrying a mutation in known breast cancer associated genes, for both epithelial and stromal compartment. **(c,d)** Scatter plots of cumulative mutation burden in relation to age, represented as number of mutations in all driver genes, per sample, in **(c)** the epithelium and **(d)** the stroma. For both plots, n = 13 parous, n = 12 nulliparous donors (some dots overlapping) and Pearson correlation (*r*) with age and P values (*P*) from linear regression are shown. NP = nulliparous, P = parous. **(e,f)** Variant allele frequency of unique recognized driver events in different samples in the **(e)** epithelial and **(f)** stromal compartment. The mutations shown in the graphs are: *TP53* (p.R273H), *KRAS* (pG12V), *PIK3CA* (p.H1047R), *PTEN* (p.R130G).

Our data, taken together with the abovementioned studies, indicates a degree of tissue specificity in the accumulation of driver events and a level of positive selection of the same genes, which will then result in further accumulation and expansion with age.

In fact, accumulation of mutations in BC-associated genes significantly increases in number with age in the nulliparous group (*P* = 1.1×10^-2^ for the totality of mutations), but not in the parous group (**Fig. 2c**). This trend was not observed in the stromal compartment, where, albeit not at significant levels, parous individuals tend to carry a higher number of mutations in the stromal cells compared to the nulliparous samples (**Fig. 2d**).

A significant increase of heterogeneity, described here as the number of mutated driver genes in each sample, was seen according to age but not parity in the epithelial compartment of the healthy breast (*P* = 0.3×10^-2^) (**Extended Data Fig. 3**).

### Recognized breast cancer driver events are present in the healthy breast

A previous study has identified not only which genes are mostly associated with BC (driver genes), but also which particular mutations may affect cancer onset or progression^23^. We confirmed the presence of 3 driver events in the healthy breast (**Fig. 2e,f**), without association of parity, albeit with one relevant exception. The missense mutation in *PTEN* resulting in p.R130Q is a recognized driver event. We recorded a different variant of this mutation in the same genomic position, resulting in p.R130G, a pathogenic variant (FATHMM score 0.96) typically associated with endometrioid carcinoma, amongst other pathologies^25^. This mutation was found at an unexpected high frequency in the healthy breast, in the epithelium of 3/25 individuals (~10%), and in the stroma of unmatched 3/25 individuals (~10%). While we filtered VAF values to ensure lack of contamination of epithelial cells, this mutation is nearly clonal (VAF >0.70) in one epithelial and one stromal sample, independently.

Known driver events in *KRAS* (pG12V) and *TP53* (p.R273H) were also found in both distinct epithelial and stromal compartment in at least 2 individuals for each group, while a mutation in *PIK3CA* (p.H1047R) was found in 2/25 stromal samples.

### Mutational signatures of the healthy breast

We observed a parity-associated increase of the percentage of T>G substitutions with age in the epithelium (*P* = 2.0×10^-2^) and both a parity-associated increase of the percentage of C>G (*P* = 2.6×10^-2^) and, to a lesser extent, a nulliparity-associated decrease of the percentage of T>C substitutions (*P* = 2.5×10^-2^) with age in the stroma (**Fig. 3a-e**). Mutations resulting in T>G and in C>G have been linked to oxidative damage^26^ and double-strand break repairs^27^, respectively. It is known that oxidative stress increases during post-partum involution in mammals^28^ and in cultured human mammary cells^29^, so we hypothesized this may be a reason for our findings.

**Figure 3.**
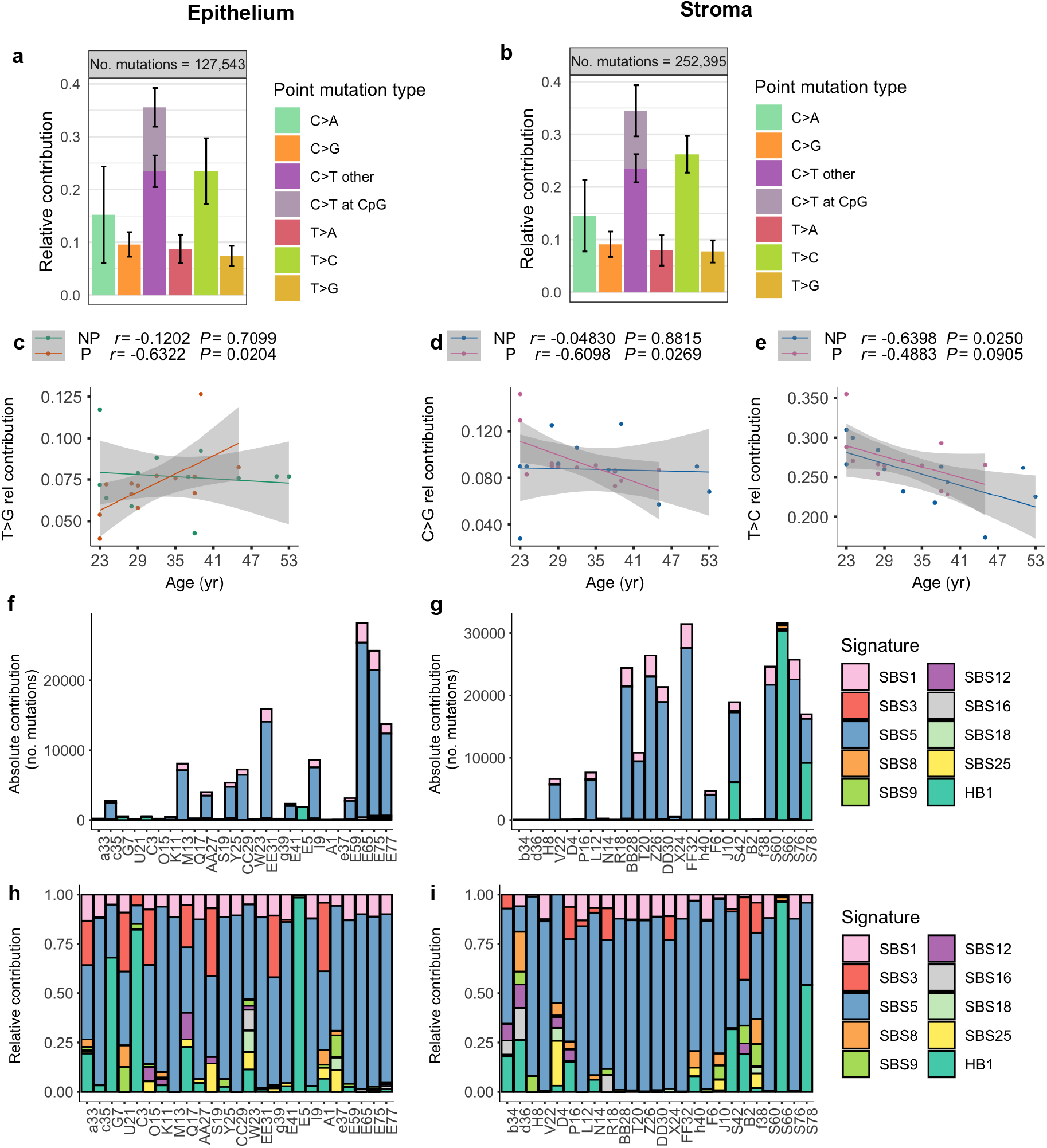
Characteristics of somatic mutations acquired in human healthy breast. **(a,b)** Relative contribution of different mutation types to the mutation spectrum for the epithelial and stromal compartment for all samples (n=25). Error bars indicate the standard deviation. **(c-e)** Scatter plot representing the correlation between the relative contribution of selected mutation types with age and pregnancy (n = 13 parous, n = 12 nulliparous donors, some dots overlapping) in the **(c)** epithelium and **(d,e)** stroma. Only mutation types with significant trends in at least one parity group are shown. Pearson correlation (*r*) with age and P values (*P*) from linear regression are shown are shown for both nulliparous (NP) and parous (P) samples. **(f-g)** Absolute and **(h-i)** relative contribution of each mutational signature from the COSMIC database and *de novo* mutation HB1 for each sample, divided in **(f,h)** epithelial and **(h,i)** stromal compartment. Samples are shown in the x axis and sorted by age of collection.

Both cellular components of our samples fitted with known single-base-substitution SBS, in particular SBS5 (unknown etiology), SBS1 (due to deamination of 5-methylcytosine) and SBS3 (associated with *BRCA1* and *BRCA2* mutations and *BRCA1* promoter methylation) (**Fig. 3f-g**). It is noteworthy that our individuals do not carry any non-synonymous mutations in either *BRCA* genes, with the exception of one individual carrying a missense mutation of uncertain significance in *BRCA2* (p.D935N, rs28897716). We did not find a specific signature associated with parity.

In the epithelium, but not in the stroma, SBS1 is a signature that was associated with age (*P* = 1.2×10^-3^, **Extended Data Fig. 4a-d**, data for stroma not shown). In the epithelium, significant increase with age was also detected for the number of occurrences constituting the following signatures: SBS5 (*P* = 1.2×10^-3^), SBS12 (*P* = 1.4×10^-3^) and SBS16 (*P* = 0.6×10^-3^). For these signatures, a significant increase with age was only seen cumulatively amongst all the samples or in the association with nulliparity (**Extended Data Fig. 4a-d**).

Importantly, the relative contribution of the signature did not significantly increase with age nor parity. The only exception to this is the relative contribution of SBS5 in the parous breast, which significantly also increased with age (*P* = 3.4×10^-2^). Several samples could not be fully categorized in any recognized signature, partly attributed to the low number of mutations of some of these samples (**Extended Data Fig. 4e and Extended Data Fig. 5a**), which would impede the construction of a mutational profile^30^. Therefore with a non-negative matrix factorization (NMF) approach, we identified a *de novo* signature, which we named HB1 (healthy breast 1) that had lower cosine similarity (< 0.9) to any individual reference cancer signatures and is characterized mostly by C>A and C>T substitutions (**Extended Data Fig. 4g-i** and **Extended Data Fig. 5b-d**).

### Gene ontology enrichment analysis of mutated genes

To determine if the healthy breast preferentially mutates in specific pathways, we selected mutations occurring in more than one sample, and analyzed if the affected genes could be enriched in specific pathways or functional groups (**Extended Data Fig. 6 and Supplementary Table 5**).

We observed several enrichment categories in the stroma. Of particular interest, mutations within the 3’ UTR can have a major impact on gene expression, affecting regions where miRNAs are binding and/or interfering with mRNA stability. Exclusively in the parous stroma, the group of genes mutated in multiple samples in the 3’ UTR were significantly enriched in pathways with transferase (GO:0016772), phosphotransferase (GO:0016773) and kinase (GO:0016301) activities, (Bonferroni q-value=2.225×10^-2^, 2.263×10^-2^ and 2.3×10^-2^, respectively, **Extended Data Fig. 6a,b**). Also, the parous stroma was characterized by an enrichment of mutated genes (missense mutations) in pathways involved with cytoskeletal organization (GO: 0007010, Bonferroni q=4.983×10^-2^, **Extended Data Fig. 6c**).

The environment of the pregnant and post-partum breast undergoes extensive remodeling of the mammary architecture. In aggregate, these data suggest that this process could be implemented thanks to the preferential accumulation of these two types of mutations.

In the epithelium, we only recorded parity-associated differences between the enrichment in genes mutated in the 5’ flank regions, where genes involved in unsaturated fatty acid metabolic processes (GO:0033559) were enriched within the nulliparous group (Bonferroni q-value =1.552×10^-2^, **Extended Data Fig. 6d**).

## DISCUSSION

Parity and age at first pregnancy are the most complex factors in BC risk, but their individual and combined contributions to the accumulation of somatic mutations in the healthy breast have not been previously investigated. In line with other healthy organs^1–3,5–10^, the mutational burden of the healthy breast significantly accumulates with age, the significance of which is particularly evident in the epithelial compartment. The absolute number of mutations is comparable with other LCM-based sequencing normal epithelia^7^. Published data derived from WGS of bulk breast tumors^23^ suggest a minimum of a 2-fold decrease in the number of substitutions in the healthy breast; however, differences in sequencing methods do not allow an exact comparison between the datasets.

The analysis of the variant allele frequencies can explain the difference in BC risk associated with age of pregnancy. Our data suggest that cell clones containing a pre-existing mutation (such as in a driver gene) in the nulliparous breast may undergo a rapid expansion during parity. Also, clone size increases with age in the parous group, suggesting that parity is an additional variable to age in the direct expansion and in the indirect probability of gaining a second driver mutation within the same cells. The combination of parity-induced and age-related clone expansion can also explain the age-independent high risk of developing breast cancer during pregnancy, in women already carrying mutant clones before childbirth. A simplified model diagram explaining the process of clonal expansion of mutated cells (one hit) can be seen in **Fig. 4**.

**Figure 4.**
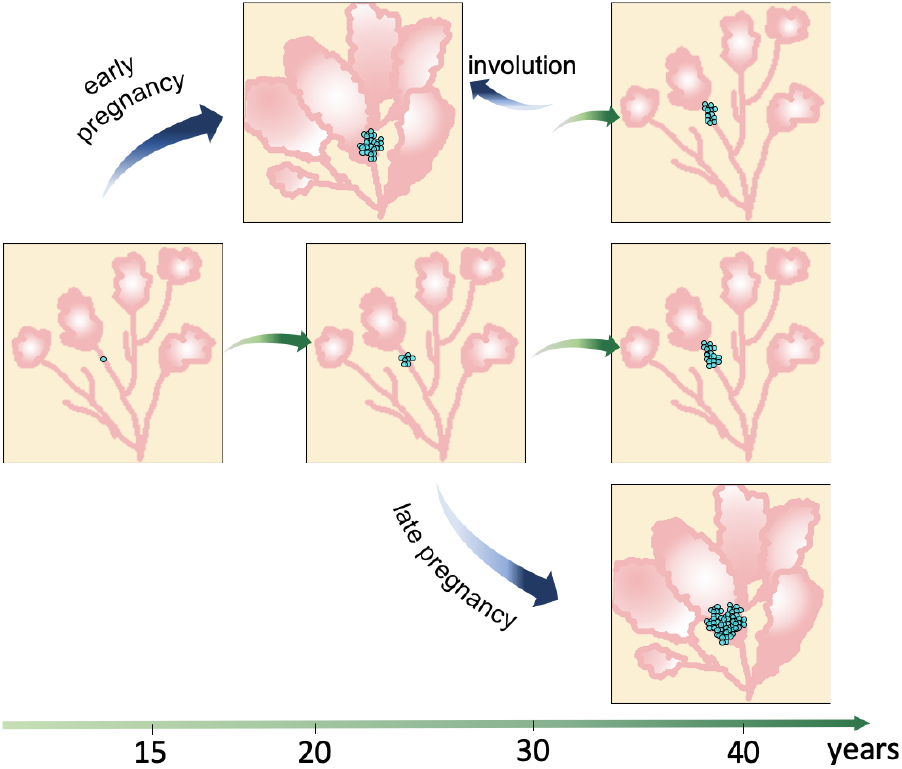
Model of expansion of a single mutated clone in the healthy breast. The scheme simplifies the probability of a clone, mutated with a single pre-existing mutation (light blue dots), to expand during age and pregnancy in the mammary epithelium (pink). Pregnancy-associated clone expansions or regression are represented by blue arrows; age-related clone expansion is represented by green arrows. In the event of life-long nulliparity (central panel), the mutated clone will expand with age, on a rate which is dependent on the mutation fitness. In the event of an early pregnancy (upper panel), the clone will be undergoing significant expansion, followed by shrinkage due to post-partum involution. The remaining clone will then keep growing at a slower rate in the breast until, possibly, a new wave of shrinkage will occur during lobular involution (not shown). In the event of a late pregnancy (bottom panel), the clone, already expanded during the years, will be undergoing even more significant expansion. However, lobular involution may not be able to significantly reduce the size of this clone, which could then progress in colonizing the epithelium.

Determining the genes that undergo positive selection in the healthy breast would provide further strength to our hypothesis and highlight functional genes which play a role in the post-pregnancy gland. To perform a confident estimate of the dN/dS ratio, we would need to use a multi-sampling approach cohort^31^, or a deeper sequencing methods, which is problematic when working with low amount of epithelial cells. However, analysis of our relatively small cohort allowed the identification of two genes possibly undergoing positive selection in the parous stroma, Sialic acid-binding Ig-like lectin family like 1 (*SIGLECL1*, q= 1.45’10^-3^) and F-Box Protein 41 (*FBXO41*, q= 0.48’10^-2^), which could be of further interest. Our current dataset did not provide evidence of positive selection in the epithelium or in the nulliparous breast.

Epidemiologic studies have determined the age of 35 as the critical age after which the protective role of pregnancy towards BC diminishes. However, our data suggests a critical shift may be closer to 32. This difference may be explained by the fact that our samples were collected from a minimum of 2 years after parturition, so the ongoing mammary involution may decrease the number of observed mutations accumulated doing pregnancy. A practical consideration for any type of genetic analysis of the mammary gland is the acknowledgment of the confounding contribution of the time passed between the year of pregnancy and the year of tissue resection to the accumulation of mutations. This information is often missing from public databases and could lead to alternative estimates. We tested the addition of 4 patients, of which the years passed between their pregnancy and the time of tissue donation was >10 years (ranging approximately from 17 to 54 years) in our analysis. Even with the limitation of the small sample size for the longer time period, it is apparent how the epithelium of the healthy mothers accumulates mutations in the years after a pregnancy (*P* = 4.7×10^-2^ for the totality of mutations) (**Extended Data Fig. 7**).

In conclusion, we believe that the mutational landscape of the healthy breast can be used as a control for the genetic study of diseases of the mammary gland, including non-malignant lesions. In BC studies, there is a need for a real “pool of normal” controls, rather than controls derived from areas adjacent or surrounding cancer, or from the contralateral breast, which have been used until now for this scope. These controls, collected from affected patients, may in fact include, albeit at low frequency, mutations which are derived from the cancer^32^. Furthermore, although BC generates from the epithelial cells of the duct, sequencing of the stromal compartment, which has not been undertaken before, may provide some insight into the elusive onset of inflammatory breast cancer, a subset of BC characterized by the dysregulation of several major inflammatory signaling pathways^33^.

## Supporting information

Key Resources Table

R markdown 1

R markdown 2 drivers

## METHODS

### Human breast samples

Twenty-nine frozen normal breast tissue samples from patients not affected by cancer were obtained from the Susan G. Komen Tissue Bank at IU Simon Cancer Center, following patient consent and approval from the local research ethics committee (IRB Protocol number is 1011003097). See **Supplementary Table 1** for clinical information. Out of these, 4 individuals were analyzed as a subgroup, represented in Supplementary Figure 7.

To confirm the normal histology, hematoxylin and eosin (H&E) staining was performed on the first section from each sample. Each section was cut at 5μm thickness, hydrated in decreasing concentration of ethanol and stained using Gills hematoxylin and 1% Eosin, as per standard protocol.

### Enzyme histochemistry

To allow visualization of the epithelium, an enzyme histochemistry targeted at a mitochondrial enzyme was performed on the section. Cytochrome C oxidase (CCO) enzyme histochemistry was performed on 16μm thick serial sections cut onto PALM membrane slides (MembraneSlide 0.17 PEN, Zeiss, Munich, Germany), as previously described^34^. Briefly, thawed sections were incubated in cytochrome c medium (100mM cytochrome c, 4mM diaminobenzidine tetrahydrochloride, 20μg/ml catalase in 0.2M phosphate buffer, pH 7.0; all from Sigma Aldrich, Poole, UK) for approximately 50min at 37°C or until satisfactory visualization of the brown color, followed by washes in PBS, pH7.4, for 3 × 5min. Sections were allowed to dry in air before proceeding to microdissection. CCO-wild type cells, present in the majority of the normal epithelium, are visible in brown, in contrast with the surrounding colorless stroma.

### Laser-capture microdissection and DNA extraction

Stained areas from the mammary epithelium, containing mammary ducts and terminal duct lobular units (TDLUs), were microdissected on a laser capture system (Zeiss PALM Microbeam) at a uniform laser power and cutting width. Approximately 30 sections were dissected for each individual. Surrounding stromal tissue with absence of detectable epithelium was superficially cut with the laser and successively isolated under a macrodissecting microscope (Leica Microsystem, Germany). Dissection and DNA extraction of the stroma and epithelium of each individual sample was performed at separate times to avoid contamination.

Pre and post dissection images were captured using a digital slide scanner and viewed using the integrated viewer software (NanoZoomer SQ, Hamamatsu Photonic, Shizuoka, Japan).

DNA was extracted using QIAamp DNA Micro kits (Qiagen, Hilden, Germany) and measured on Qubit 3.0 Fluorimeter, using Qubit dsDNA HS Assay kit (Life Technologies, Paisley, UK), according to the manufacturer’s recommendation.

### Library preparation and DNA sequencing

Library preparation and sequencing was carried out by the Beijing Genomic Institute (Shenzhen, Guangdong, China). In brief, isolated DNA (100ng) was fragmented by Covaris technology to obtain fragments of an average 350bp length. The repaired/dA tailed DNA fragments were then ligated to both ends with the adapters and amplified by ligation-mediated PCR (LM-PCR), followed by single strand separation and cyclization. A rolling circle amplification (RCA) was performed to produce DNA Nanoballs (DNBs). The qualified DNBs were loaded into patterned nanoarrays and 100bp pair-end read sequences were read on the BGISEQ-500 platform, which studies have shown to be of comparable sensitivity to other commercial platforms^35^.

### Mutations calling and downstream analysis

Alignment of the reads and calling were carried out by the bioinformatic team at the Beijing Genomic Institute, following the guidelines of the Broad Institute Genome Analysis Toolkit (GATK, https://www.broadinstitute.org/gatk/guide/best-practices). Briefly, cleaned data for each individual sample was mapped to the human reference genome (GRCh37/HG19) with Burrows-Wheeler Aligner (v0.7.12) and duplicated reads were removed by Picard-tools (v1.118) with default settings. Details of quality control, coverage and mapping can be found in **Supplementary Table 2**.

Somatic short mutations, including SNV, small insertions and deletions, were called via local assembly of haplotypes with Mutect2 (v4.1.4.1, https://gatk.broadinstitute.org/hc/en-us/sections/360007458971-4-1-4-1) with an adaptation of the “tumor with matched normal” function, where tumor was substituted with normal epithelium, and matched normal was substituted with matched normal stroma. Default settings for tumor-normal pairs were used. Called variants were successively filtered with FilterMutectCalls (v4.1.4.1, https://gatk.broadinstitute.org/hc/en-us/sections/360007458971-4-1-4-1), and only mutations categorized with PASS, representing confidently called somatic mutations, were used in the downstream analysis. VCF files were converted to Mutation Annotation Format (MAF) files according to National Cancer Institute specifications using Mskcc/vcf2maf (v1.6.17)^36^, adapting the format to include epithelial and stromal barcodes instead of tumor and normal barcodes, respectively.

Annotated mutations were then further filtered for (1) total number of reads in both cellular compartment > 10; (2) mutated number of reads > 5; variant allele frequency (VAF) >0.02%; (3) frequency > 0.1% in the Genome Aggregation database (gnomAD, V2.1.1, exome and genome samples, https://gnomad.broadinstitute.org/), and in the 1000 Genome database (https://www.internationalgenome.org/). MAF were visualized and summarized using the R Bioconductor package, Maftools (v1.8.10)^37^ when necessary before proceeding to detailed analysis. Variants were categorized into synonymous and non-synonymous according to the impact on protein coding of each variants (Ensembl IMPACT rating). Non-synonymous variants with high or moderate IMPACT were: Frame shift deletions, Frame shift insertions, Nonsense mutations, Nonstop mutations, Splice sites, Translation start sites, In frame deletions, In Frame Insertions, Missense mutations. Synonymous mutations with low or modifier IMPACT were: 3’ Flank, 5’ Flank, 3’ UTR, 5’ UTR, Intron, IGR, Splice regions, RNA, Silent mutations.

Downstream analysis was conducted in R^38^ and all figures were produced using the package ggplot2(v3.3.0)^39^, unless otherwise specified.

### Determination of mutations occurring in breast cancer-associated genes

The list of identified breast cancer drivers which we have included in our analysis were taken from the paper “Landscape of somatic mutations in 560 breast cancer whole-genome sequences” by Nik-Zainal *et al*., Nature, 2016^23^. Individual driver events were taken from the same manuscript and found in Supplementary Table 14: Driver events by mutation type 01052015.v2.

### Detection of positively selected genes

To determine dN/dS (calculated on the observed/expected ratios of non-synonymous to synonymous mutations), we combined all the somatic mutations from the 29 individuals, and used the default settings of dNdScv (version 0.0.1.0)^40^, with significant genes determined at q<0.01.

### Identification of mutational signatures

Mutational signatures were identified using the R Bioconductor package MutationalPatterns (v2.0.0)^30^. Briefly, VCF files were imported as *GRanges* object and the sequence context was derived from the imported Reference Genome hg19, installed with the R Bioconductor package BSGenome (v1.56.0)^41^. The contribution of known signatures from the COSMIC database^25^ was determined, and, for samples of adequate number of mutations but low cosine similarity with any known signature, two *de novo* signatures were extracted using non-negative matrix factorization (NMF). Mutational profile similarity was then evaluated to compare the two signatures with the ones from the COSMIC database, and to allow the inclusion of only one *de novo* signature, which we named HB1 (Healthy Breast 1).

### Gene ontology analysis

Each functional category was analyzed using the ToppGene portal^42^ (https://toppgene.cchmc.org/) to identify enrichments in Gene Ontology (GO) and pathways. Each p-value was corrected for multiple comparison with Bonferroni correction and shown in the graph as the inverse of log10 of the adjusted p-value (-log10(q)) for easiness of interpretation. Significant outputs were filtered out if each dataset contained less than 10 genes for group. Categories included in these studies were gene ontology (GO) annotations (biological process, cellular component and molecular function), and pathway annotations (Kyoto Encyclopedia of Genes and Genomes, KEGG).

## DATA AVAILABILITY

All sequencing data and accession codes will be made available via the European Genome-Phenome Archive prior to publication (https://ega-archive.org/).

## CODE AVAILABILITY

We used published software and algorithms for this study. **See Key Resource Table**. Code for statistical analyses on total substitution and driver mutation burdens is included in the Supplementary Information. All other code is available on request.

## AUTHOR CONTRIBUTIONS

BC and JS designed the study, analyzed data and wrote the manuscript. NT performed LCM, DNA extraction and staining on all samples. LDDB, AJJ contributed to experimental work and to data analysis. Contribution on data filtering for somatic mutations (CM), mutational analysis of the stroma (SZ), bioinformatic support, data analysis and discussion (AY, BW and NM). All authors reviewed, contributed to and approved the submitted version.

## ACKNOWLEDGMENTS

BC, NT, DBBL, SZ and NM are funded by Action Against Cancer and UKRI-IBIN (NM). AY is funded by British Heart Foundation. BW is funded by a Barts Charity Lectureship (grant MGU045).

Samples from the Susan G. Komen Tissue Bank at the IU Simon Cancer Center were used in this study. We thank contributors, including Indiana University who collected samples used in this study, as well as donors and their families, whose help and participation made this work possible.

We would like to thank Dr Linda Hammond (Microscopy Facility, Barts Cancer Institute, Queen Mary University of London), Prof Rob Krams (Dept of Bioengineering, Imperial College London and currently at Queen Mary University of London), Prof Jesus Gil (MRC London Institute of Medical Sciences) and Prof Simone di Giovanni (Dept of Brain Science, Imperial College London) for the kind use of the laser-capture microscopes, and Dr Veronique Azuara (Department of Metabolism, Digestion and Reproduction, Imperial College London) for the use of the macrodissection microscope. Finally, we would like to thank the late Prof Sami Shousha (Imperial College London) for histopathology advice. We acknowledge support of the NIHR and Imperial BRC and ECMC.

## EXTENDED DATA FIGURES

**Extended Data Figure 1.**
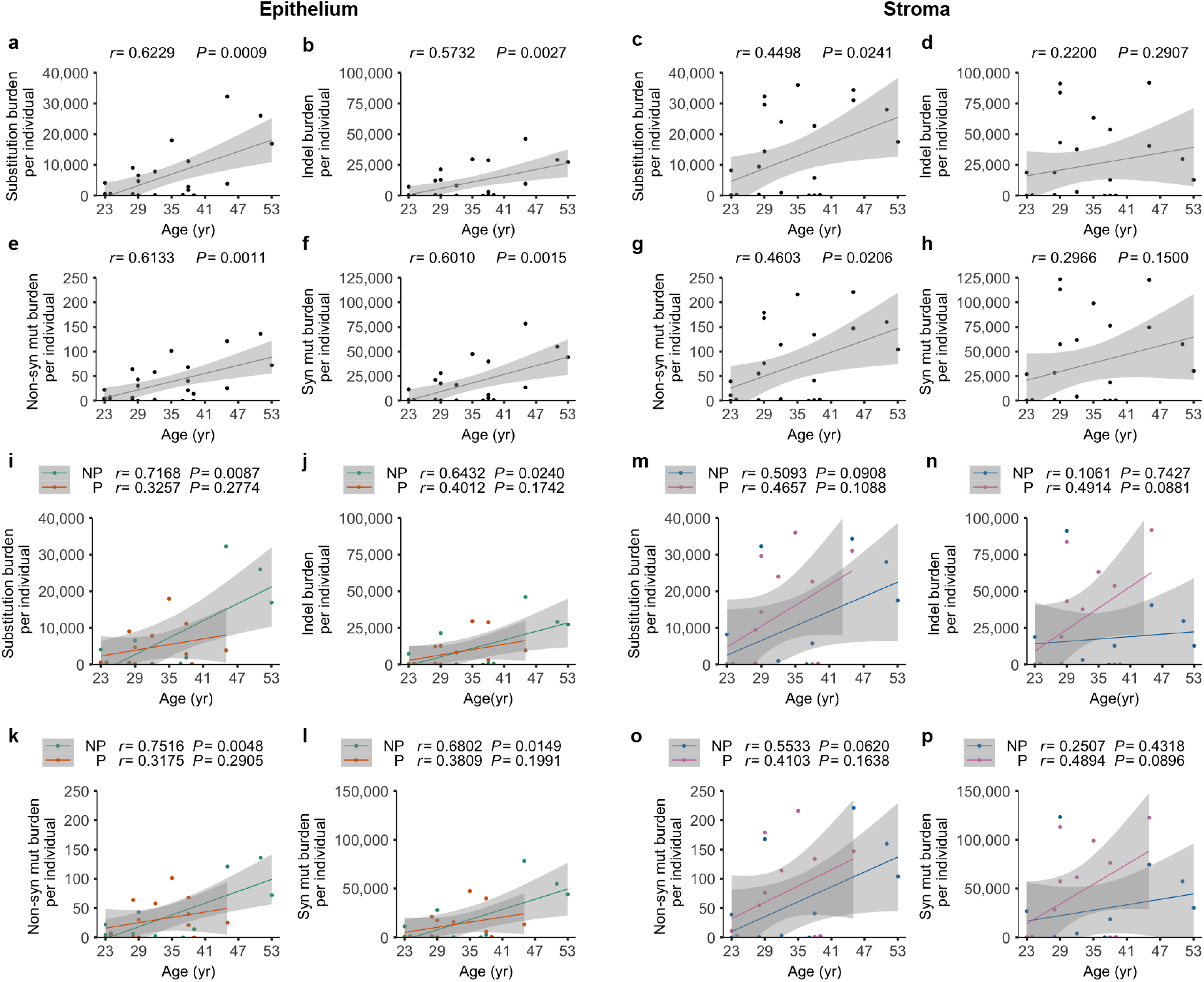
Correlation of the mutation burden with age of collection and parity in the healthy breast. **(a-h)** Mutation burden shown as one dot for each donor (n = 25 donors, some dots overlapping). Pearson correlation (*r*) with age and P values (*P*) from linear regression are shown for the epithelial (left) and stromal (right) compartment. **(a-h)** Correlation between substitutions, indel, non-synonymous and synonymous burden with age is shown. **(i-p)** correlation between substitutions, indel, non-synonymous and synonymous burden with age and pregnancy is shown. For parity plots, n = 13 parous, n = 12 nulliparous donors (some dots overlapping) and Pearson correlation (*r*) with age and P values (*P*) from linear regression are shown. NP = nulliparous, P = parous.

**Extended Data Figure 2.**
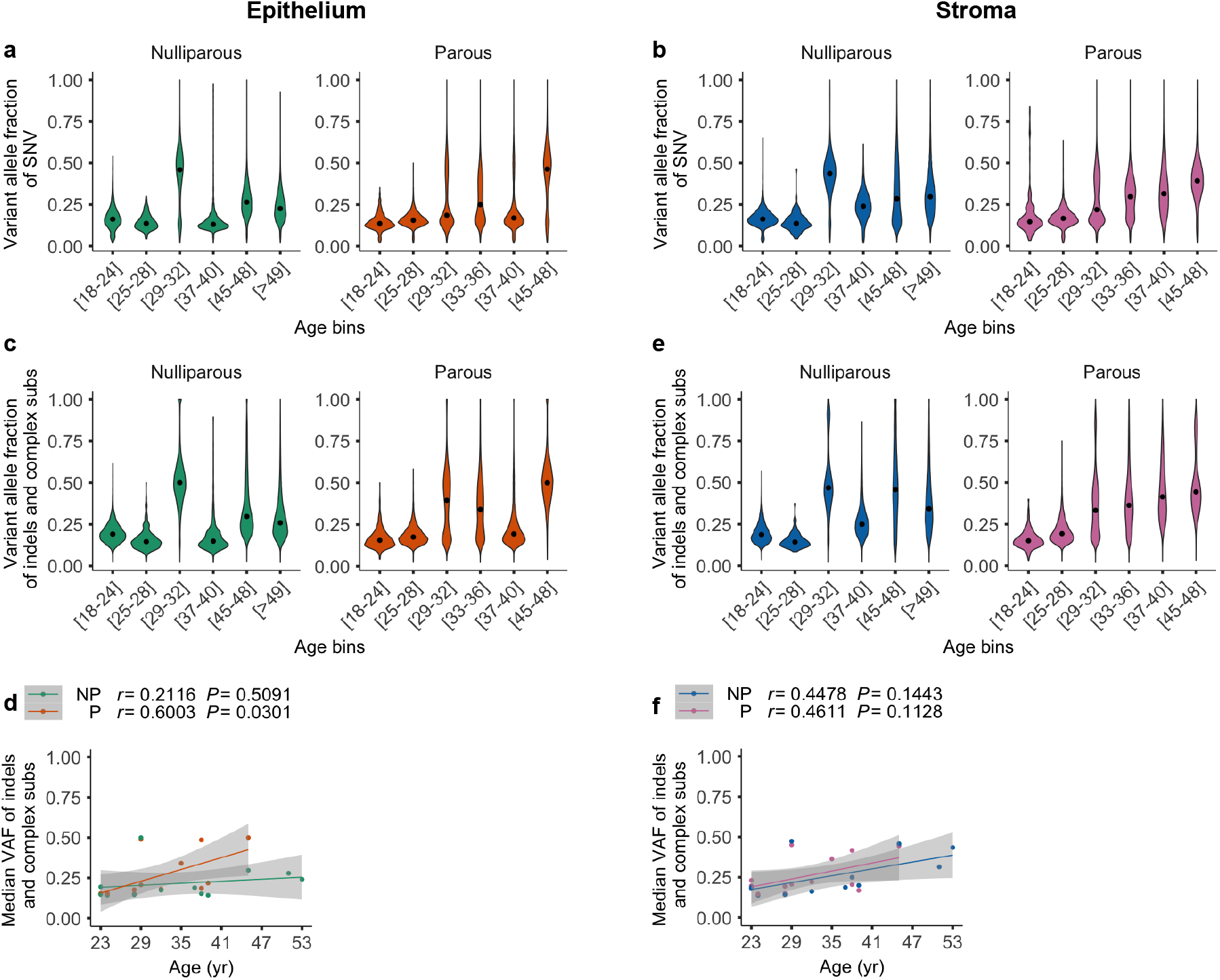
Total mutation burden and clonality in the healthy parous and nulliparous breast. **(a,e)** Violin plots of variant allele frequencies of **(a,b)** SNV and **(c,e)** indels and complex substitutions for each parity group for the epithelial (left) and stromal (right) compartment. Samples are binned in regular 4-years age intervals. Black dots represent median values. The bigger variation in the [29,32] age range is due to a particularly mutated individual **(d,f)** Plots of median variant allele frequency of indels and complex substitutions for each parity group. For both plots, n = 13 parous, n = 12 nulliparous donors (some dots overlapping) and Pearson correlation (*r*) with age and P values (*P*) from linear regression are shown. NP = nulliparous, P = parous.

**Extended Data Figure 3.**
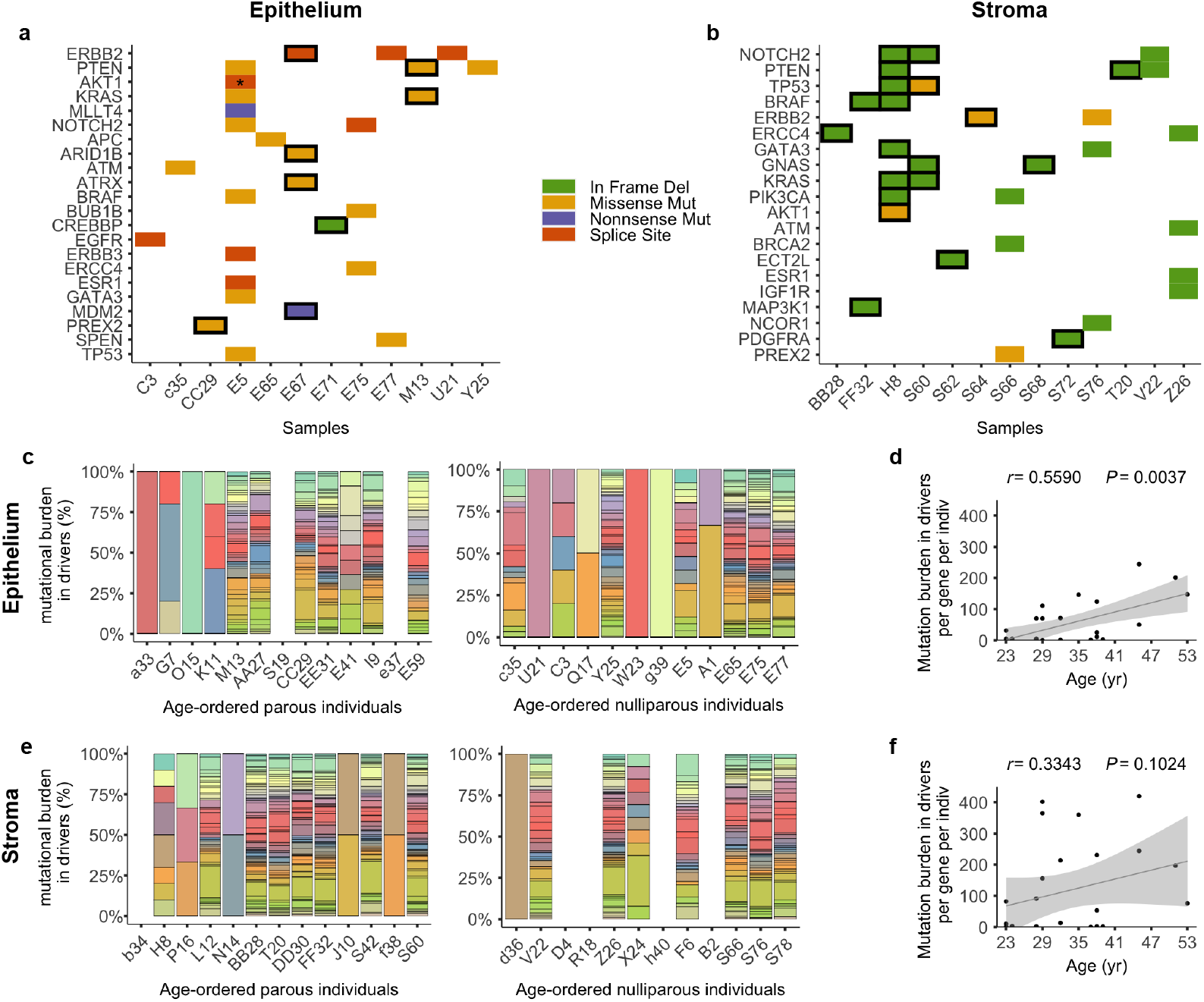
Heterogeneity of driver genes in the healthy breast. **(a,b)** Number of non-synonymous mutations in breast cancer driver genes per individual for the **(a)** epithelial and **(b)** stromal compartment. Every tile represents one occurrence, with the exception of *AKT1* (asterisk), which contained 2 splice site mutations. Tiles bordered in black indicate parous individuals. **(c,e)** Heterogeneity in the mutational burden in driver genes is shown by samples for epithelium and stroma. Each color represents a distinct mutated driver gene. Samples without colored bar indicate lack of mutations in driver genes. **(d,f)** Scatter plot of mean mutational burden per gene per individual, for both **(c)** epithelium and **(d)** stroma. For both plots, n = 25 donors (some dots overlapping) and Pearson correlation (*r*) with age and P values (*P*) from linear regression are shown.

**Extended Data Figure 4.**
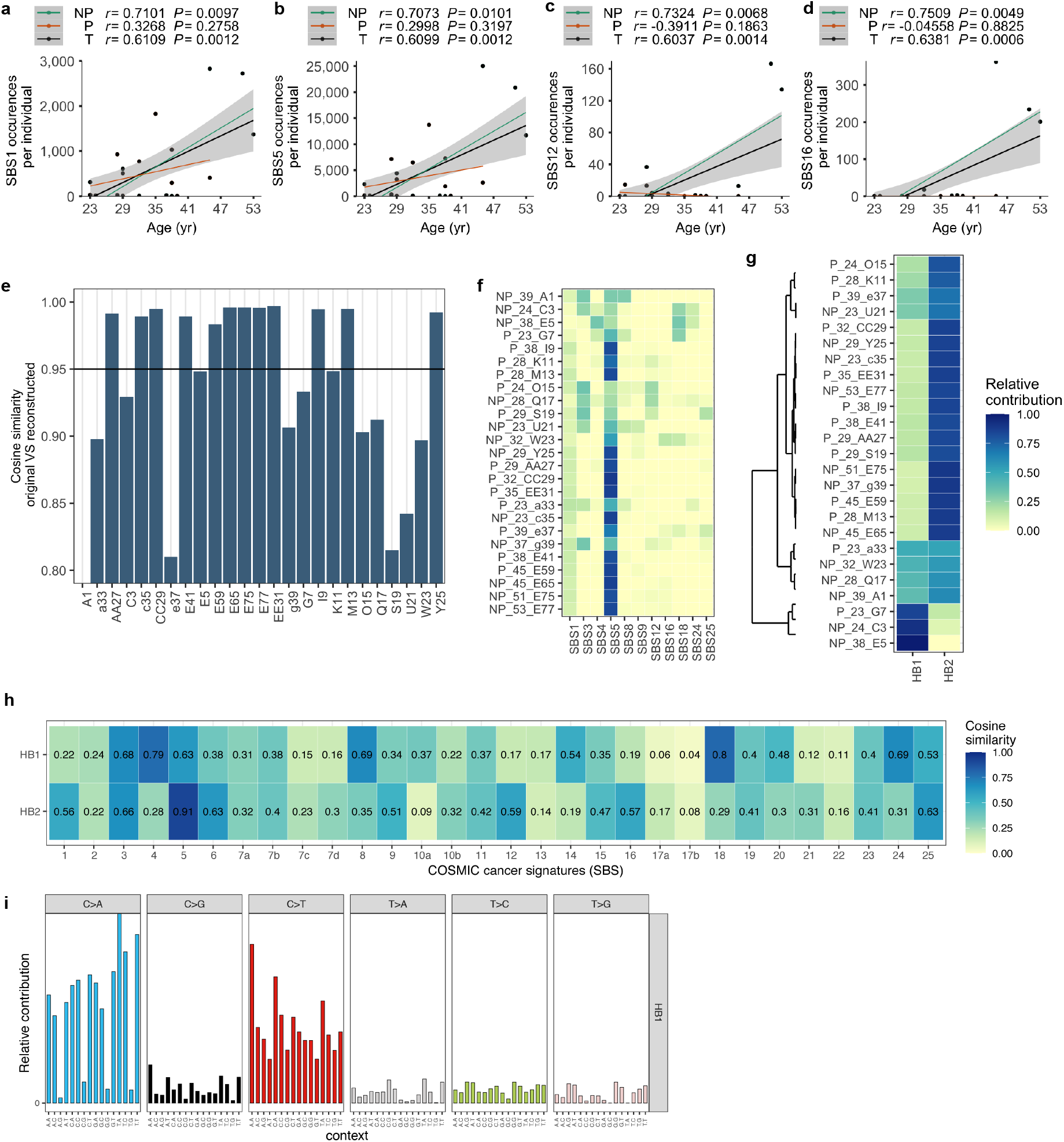
Mutational signatures in the epithelium of the healthy breast. **(a,d)** Scatter plot showing correlations with age for the mutation burdens attributable in each parity group (NP, nulliparous, n=12; P, parous, n=15; T, combined samples) for SBS1, SBS5, SBS12, SBS16. **(e-h)** Reconstruction of mutational profiles using known COSMIC and *de novo* identified signatures. **(e)** Cosine similarity between the original mutational profile and the reconstructed mutational profile based on the optimal linear combination of all COSMIC v3.1 signatures for each sample. Threshold of cosine similarity is shown by the horizontal line. Samples are not shown in a particular order. **(f)** Heatmap showing the optimal relative contribution of COSMIC signatures v3.1 in each sample. Only signatures with at least 10% contribution in at least one of the samples are shown. Samples are shown in the format “parity_age_name”. **(g)** Heatmap showing the relative contribution of two mutations extracted *de novo* using NMF, HB1 and HB2, in each sample. Samples are shown in the same format as above. **(h)** Heatmap showing the cosine similarity of the extracted signatures in **(g)** with the COSMIC signatures v3.1. **(i)** Mutational profile of signature HB1 (conventional 96 mutation type classification).

**Extended Data Figure 5.**
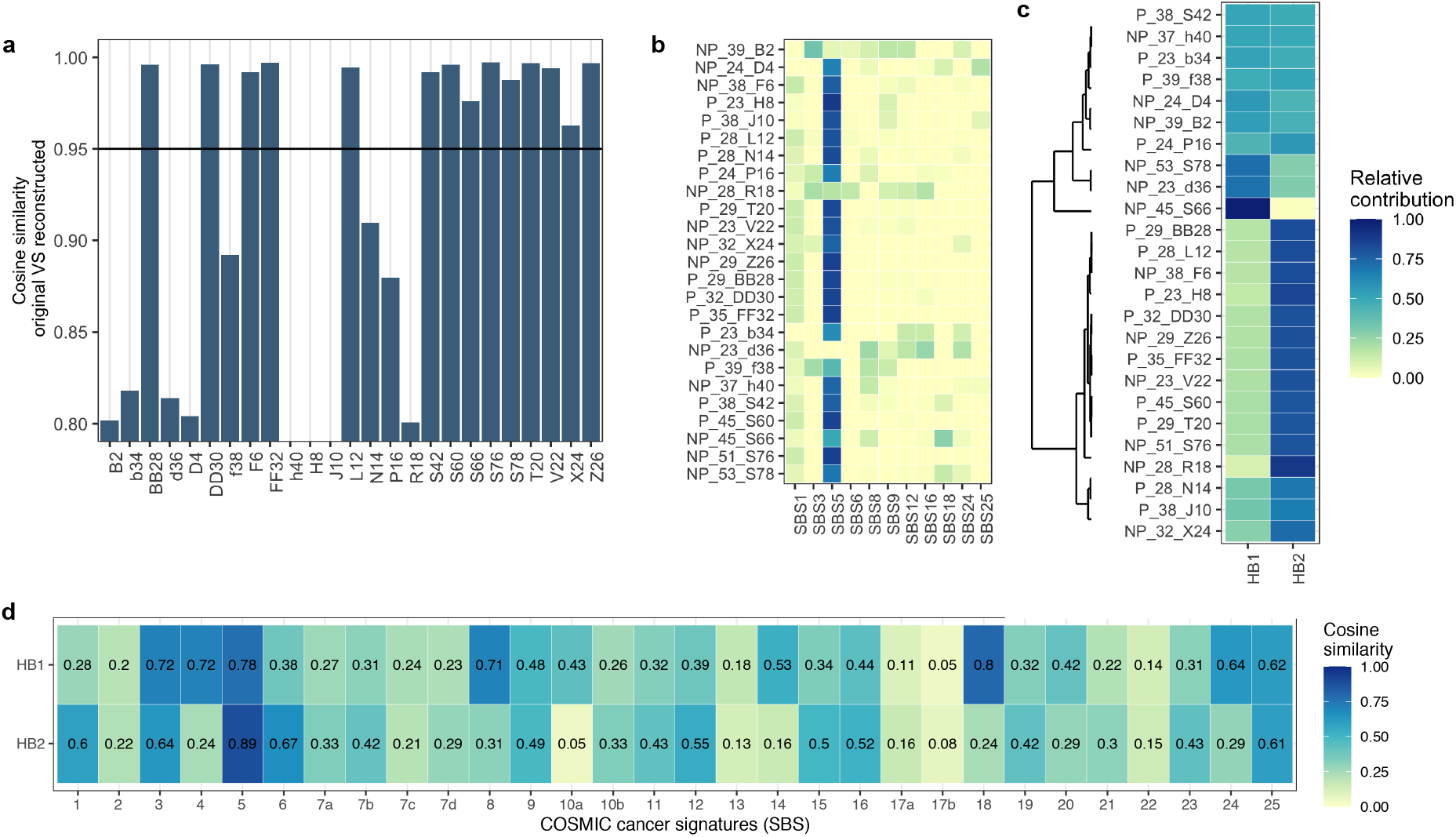
Mutational signatures in the stroma of the healthy breast. Reconstruction of mutational profiles using known COSMIC and *de novo* identified signatures. **(a)** Cosine similarity between the original mutational profile and the reconstructed mutational profile based on the optimal linear combination of all COSMIC v3.1 signatures for each sample. Threshold of cosine similarity is shown by the horizontal line. Samples are not shown in a particular order. **(b)** Heatmap showing the optimal relative contribution of COSMIC signatures v3.1 in each sample. Only signatures with at least 10% contribution in at least one of the samples are shown. Samples are shown in the format “parity_age_name”. **(c)** Heatmap showing the relative contribution of two mutations extracted *de novo* using NMF, HB1 and HB2, in each sample. Samples are shown in the same format as above. **(d)** Heatmap showing the cosine similarity of the extracted signatures in **c** with the COSMIC signatures v3.1.

**Extended Data Figure 6.**
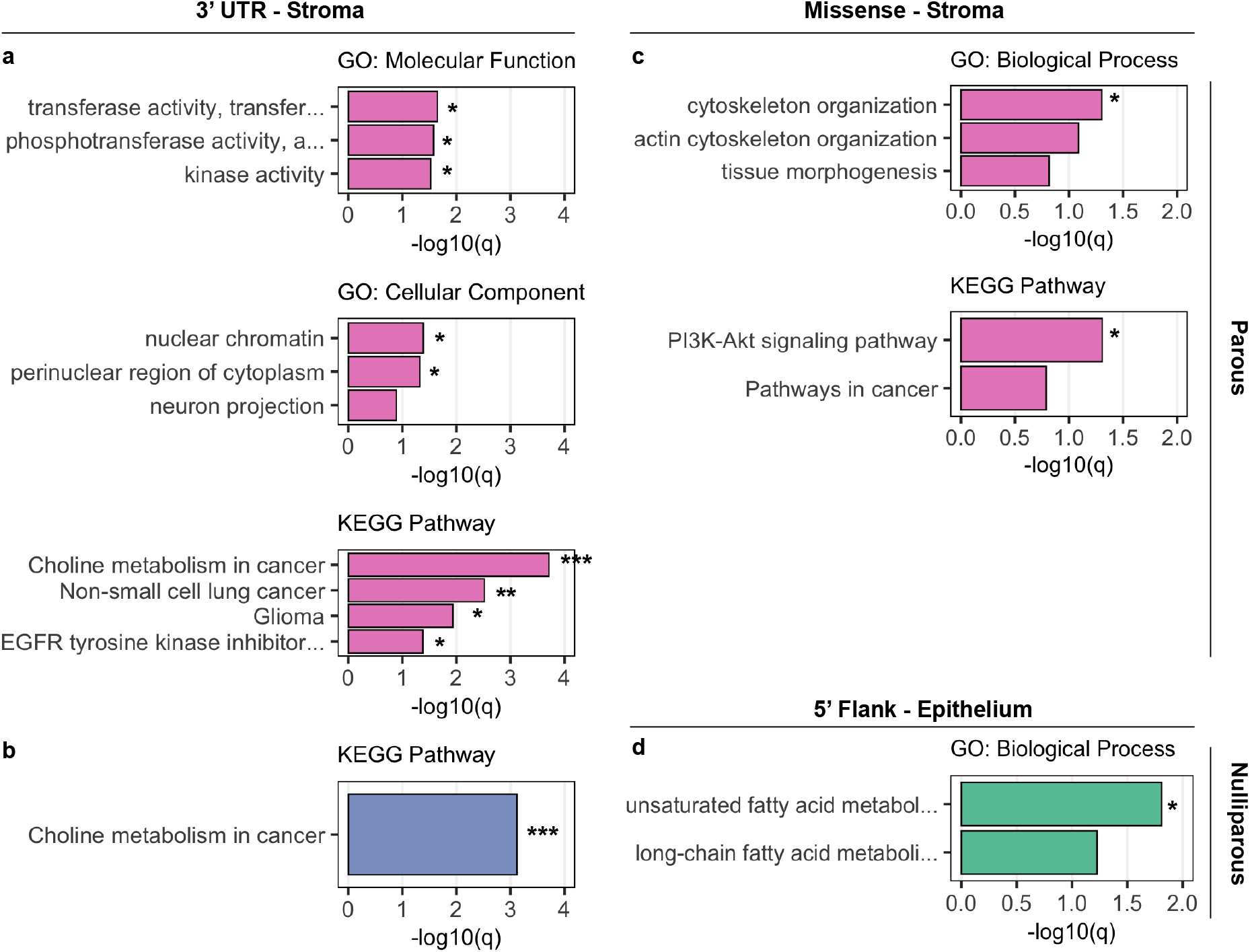
Gene ontology enrichment analysis of mutated genes. GO enrichment analysis of mutated genes was performed using ToppGene suite (see Methods). Only significantly or close to significant enriched GO terms in biological process, molecular function and cellular component categories and in KEGG pathways are shown. The Bonferroni-adjusted statistically significant values were negative 10-base log transformed. Analysis is shown for: **(a)** genes mutated in the parous stroma and carrying mutations in 3’ UTR regions; **(b**) genes mutated in the nulliparous stroma and carrying mutations in 3’ UTR regions; **(c**) genes mutated in the parous stroma and carrying missense mutations; **(d**) genes mutated in the nulliparous epithelium and carrying mutations in 5’ Flank regions. Asterisks represent significance thresholds: q<0.05 (*); q<0.01 (**); q<0.001 (***).

**Extended Data Figure 7.**
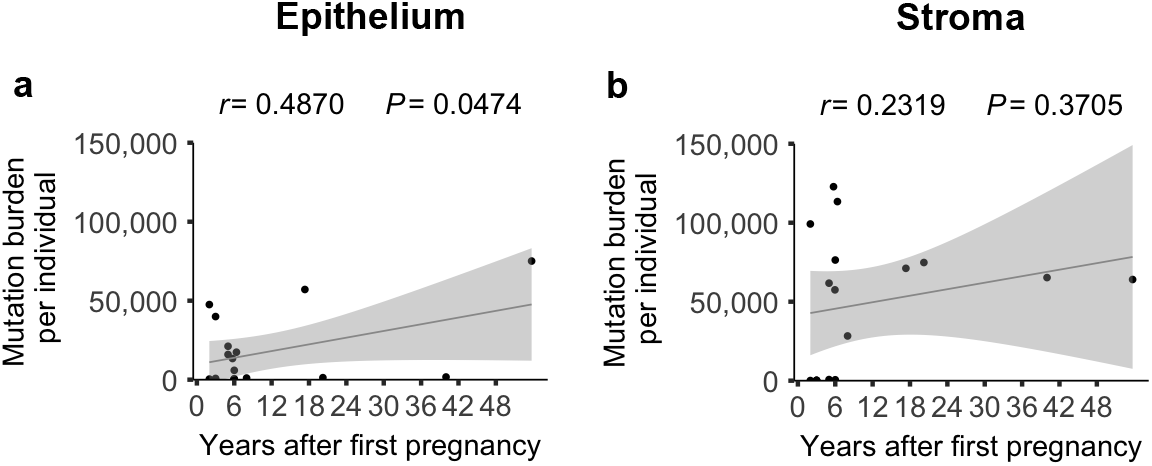
Change of the mutation burden with years after partum in the healthy parous breast. Mutation burden shown as one dot for each donor (n = 17 donors, some dots overlapping). Pearson correlation (*r*) with age and P values (*P*) from linear regression are shown are shown for the epithelial **(a)** and stromal **(b)** compartment.

