## Supplementary material for "Mutational landscapes of normal breast during age and pregnancy determine cancer risk": Key Resources Table

**The mutational landscape of the normal breast during age and pregnancy determines cancer risk**

Cereser *et al.*, 2020

### KEY RESOURCE TABLE

| REAGENT or RESOURCE | SOURCE | IDENTIFIER |
|---|---|---|
| Healthy breast Tissue Bank | Susan G. Komen Tissue Bank at IU Simon Cancer Center | https://komentissuebank.iu.edu/ |
| Mutect2 (v4.1.4.1): Algorithm for calling somatic short mutations via local assembly of haplotypes | Broad Institute of MIT and Harvard | https://gatk.broadinstitute.org/hc/en-us/articles/360037593851-Mutect2 |
| FilterMutectCalls (v4.1.4.1): Algorithm for filtering somatic SNVs and indels called by Mutect2 | Broad Institute of MIT and Harvard | https://gatk.broadinstitute.org/hc/en-us/articles/360037225412-FilterMutectCalls |
| vcf2maf (v1.6.17)): Algorithm for converting a VCF into a MAF | (Kandoth, 2019) | https://github.com/mskcc/vcf2maf |
| List of driver events from breast cancer study | (Nik-Zainal et al., 2016) | Supplementary Table 14: Driver events by mutation type 01052015.v2. |
| gnomAD, v2.1.1: collection of call sets from exome and genome sequencing data | Broad Institute of MIT and Harvard | https://gnomad.broadinstitute.org/ |
| 1000 Genome database: public catalogue of human variation and genotype data | IGSR: The International Genome Sample Resource, EMBL-EBI | https://www.internationalgenome.org/ |
| dNdScv (v0.0.1.0): R package, group of maximum-likelihood dN/dS methods to quantify selection in cancer and somatic evolution | (Martincorena et al., 2017) | https://github.com/im3sanger/dndscv |
| MutationalPatterns (v2.0.0): R package for the characterization and of mutational patterns | (Blokzijl et al., 2018) | https://bioconductor.org/packages/release/bioc/html/MutationalPatterns.html |
| BSGenome (v1.56.0): R package for representation of full genomes and their SNPs | (Pagès, 2020) | http://bioconductor.org/packages/release/bioc/html/BSgenome.html |
| ToppGene Suite: gene list enrichment analysis and candidate gene prioritization | (Chen et al., 2009) | https://toppgene.cchmc.org/ |
