## Supplementary material for "Mutational landscapes of normal breast during age and pregnancy determine cancer risk": R markdown 1

B Cereser

### Mutational burden of the normal breast during age and pregnancy

### Analysis of the mutational burden and VAF in the epithelium

##### Load Libraries

```
library(data.table)
library(ggplot2)
library(RColorBrewer)
library(scales)
library(dplyr)
```

##### Load in data files

Load in sample data for the total number of donors with associated meta-data.

```
Ep_somatic <- fread("Ep_somatic.txt.gz", sep="\t", header = T)
```

##### Analyse only either NP samples or parous (P) samples where the gap between parturition and tissue collection is < 10 years

This will leave you with the 25 samples analysed in the main paper. The excluded 4 samples are the ones analyzed in the last Extended Data Figure. The column Years_after_pregn (Years after pregnancy) represents the gap between parturition and tissue collection. For Nulliparous (NP) is artificially set at 0 to make it easier when plotting the sets together.

```
Ep_somatic_close<-subset (Ep_somatic, Years_after_pregn <10)
```

### 1. Correlation beetween age/parity on all somatic mutations of the epithelium

##### Count to determine how many mutations each sample accumulates, divided for age group

```
Ep_somatic_close$Parity = factor(Ep_somatic_close$Parity)
Ep_somatic_close$Ep_Sample_Barcode = factor(Ep_somatic_close$Ep_Sample_Barcode)
Ep_somatic_close_count<-Ep_somatic_close %>% group_by(Age, Parity,
         Ep_Sample_Barcode) %>% tally
```

##### Plot mutation burden vs age, independently of parity and calculate correlation and linear regression

```r
ggplot(Ep_somatic_close_count, aes(x= Age, y=n)) +
  geom_point(size = 0.2) +
  geom_smooth( colour = "#999999", method = 'lm', size = 0.2) +
  scale_x_continuous(breaks = round(seq(min(Ep_somatic_close_count$Age),
          max(Ep_somatic_close_count$Age), by = 6),1)) +
  scale_y_continuous(labels = comma) +
  scale_color_brewer(palette = "Dark2") +
  labs(x= "Age (yr)", y = "Mutation burden \n per individual")
```

```
## `geom_smooth()` using formula 'y ~ x'
```

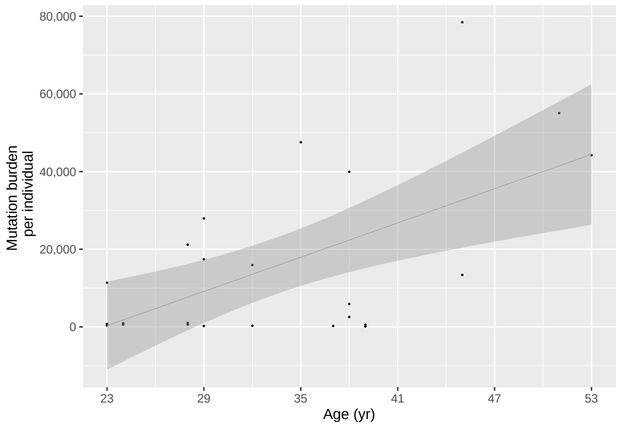


```r
cor(Ep_somatic_close_count$n, Ep_somatic_close_count$Age)
```

```
## [1] 0.6010532
```

```r
summary(lm(formula =  Ep_somatic_close_count$n ~ Ep_somatic_close_count$Age))
```

```
##
## Call:
## lm(formula = Ep_somatic_close_count$n ~ Ep_somatic_close_count$Age)
##
## Residuals:
##    Min     1Q Median     3Q    Max
## -23722 -13246   -194  11063  45783
##
## Coefficients:
##                            Estimate Std. Error t value Pr(>|t|)
## (Intercept)                -33474.1    14024.1  -2.387  0.02560 *
## Ep_somatic_close_count$Age   1469.5      407.4   3.607  0.00149 **
## ---
## Signif. codes:  0 '***' 0.001 '**' 0.01 '*' 0.05 '.' 0.1 ' ' 1
##
## Residual standard error: 17590 on 23 degrees of freedom
## Multiple R-squared:  0.3613, Adjusted R-squared:  0.3335
## F-statistic: 13.01 on 1 and 23 DF,  p-value: 0.001485
```

**Plot mutation burden vs age and parity and calculate linear regression for each parity group**

```r
Ep_somatic_P <- Ep_somatic_close[Ep_somatic_close$Parity %in% "Parous"]
Ep_somatic_NP <- Ep_somatic_close[Ep_somatic_close$Parity %in% "Nulliparous"]

Ep_somatic_P_count<-Ep_somatic_P %>% group_by(Age, Parity, Ep_Sample_Barcode) %>% tally
Ep_somatic_NP_count<-Ep_somatic_NP %>% group_by(Age, Parity, Ep_Sample_Barcode) %>% tally

#parous
cor(Ep_somatic_P_count$n , Ep_somatic_P_count$Age)
```

```
## [1] 0.3808474
```

```r
summary(lm(formula =  Ep_somatic_P_count$n ~ Ep_somatic_P_count$Age))
```

```
##
## Call:
## lm(formula = Ep_somatic_P_count$n ~ Ep_somatic_P_count$Age)
##
## Residuals:
##    Min     1Q Median     3Q    Max
## -18972 -10119  -4777   7032  31954
##
## Coefficients:
##                        Estimate Std. Error t value Pr(>|t|)
## (Intercept)            -14886.5    20599.9  -0.723    0.485
## Ep_somatic_P_count$Age    871.1      637.7   1.366    0.199
##
## Residual standard error: 15270 on 11 degrees of freedom
## Multiple R-squared:  0.145,  Adjusted R-squared:  0.06732
## F-statistic: 1.866 on 1 and 11 DF,  p-value: 0.1992
```

```r
#nulliparous
cor(Ep_somatic_NP_count$n , Ep_somatic_NP_count$Age)
```

```
## [1] 0.6804236
```

```r
summary(lm(formula =  Ep_somatic_NP_count$n ~ Ep_somatic_NP_count$Age))
```

```
##
## Call:
## lm(formula = Ep_somatic_NP_count$n ~ Ep_somatic_NP_count$Age)
##
## Residuals:
##    Min     1Q Median     3Q    Max
## -24696 -14798  -1787  10209  42765
##
## Coefficients:
##                         Estimate Std. Error t value Pr(>|t|)
## (Intercept)             -42623.5    21673.0  -1.967   0.0776 .
## Ep_somatic_NP_count$Age   1739.9      592.6   2.936   0.0149 *
## ---
## Signif. codes:  0 '***' 0.001 '**' 0.01 '*' 0.05 '.' 0.1 ' ' 1
```

```
##
## Residual standard error: 20630 on 10 degrees of freedom
## Multiple R-squared:  0.463,  Adjusted R-squared:  0.4093
## F-statistic: 8.621 on 1 and 10 DF,  p-value: 0.01488
```

```
#
ggplot(Ep_somatic_close_count, aes(x= Age, y=n, color= Parity)) +
  geom_point(size=0.2) +
  geom_smooth(aes(colour=Parity), method = 'lm', size = 0.2) +
  scale_x_continuous(breaks = round(seq(min(Ep_somatic_close_count$Age),
        max(Ep_somatic_close_count$Age), by = 6),1)) +
  scale_y_continuous(labels = comma) +
  scale_color_brewer(palette = "Dark2") +
  labs(x= "Age (yr)", y = "Mutation burden \n per individual")
```

```
## `geom_smooth()` using formula 'y ~ x'
```

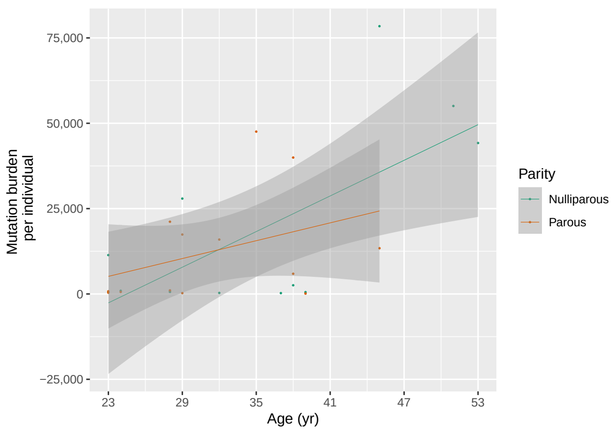


**Plots for individual substitutions, indels, synonym and non-synonym mutations were run with the same codes as above, based on the appropriate following subsets.**

```
#substitutions
subs<- c("SNP","DNP","TNP")
Ep_subs <- Ep_somatic_close[Ep_somatic_close$Variant_Type %in% subs,]

#indels
indel <- c("DEL"    ,"INS")
Ep_indel <- Ep_somatic_close[Ep_somatic_close$Variant_Type %in% indel,]

#Syn and non-syn
nonSyn <- c("Frame_Shift_Del", "Frame_Shift_Ins", "Splice_Site","Translation_Start_Site",
          "Nonsense_Mutation", "Nonstop_Mutation", "In_Frame_Del","In_Frame_Ins",
          "Missense_Mutation")
Ep_somatic_close$Class<-ifelse(Ep_somatic_close$Variant_Classification %in% nonSyn,
          'NonSyn', 'Syn')
Ep_Syn <- Ep_somatic_close[!Ep_somatic_close$Class %in% "NonSyn",]
Ep_NonSyn <- Ep_somatic_close[Ep_somatic_close$Class %in% "NonSyn",]
```

### 2. Correlation beetween VAF of SNV with age and parity

#### Extract only the SNV from the entire data

```
Ep_somatic_SNV <- Ep_somatic_close[Ep_somatic_close$Variant_Type %in% "SNP",]
```

#### Calculate the median VAF

```
Epithelial_snv_median<-aggregate(Ep_somatic_SNV[, 15],
          list(Ep_somatic_SNV$Ep_Sample_Barcode), median)
#rename the column
colnames(Epithelial_snv_median)[1]<-"Ep_Sample_Barcode"
#re-join info on Age and Parity to the median
sample_age <- fread("clin_info.txt", sep="\t", header = T)
colnames(sample_age)[1]<-"Ep_Sample_Barcode"
sample_age<-sample_age[,c(1,3,4)]
Epithelial_snv_median1<-left_join(Epithelial_snv_median, sample_age,
          by = "Ep_Sample_Barcode")
```

#### Plot median VAF of SNV with age for each parity group and calculate linear regression

```
Ep_snv_P_median <- Epithelial_snv_median1[Epithelial_snv_median1$Parity %in% "Parous",]
Ep_snv_NP_median <- Epithelial_snv_median1[Epithelial_snv_median1$Parity
          %in% "Nulliparous",]

cor_P <- cor(Ep_snv_P_median$Ep_vaf , Ep_snv_P_median$Age)
P <- summary(lm(formula =  Ep_snv_P_median$Ep_vaf ~ Ep_snv_P_median$Age))
cor_NP <- cor(Ep_snv_NP_median$Ep_vaf , Ep_snv_NP_median$Age)
NP <- summary(lm(formula =  Ep_snv_NP_median$Ep_vaf ~ Ep_snv_NP_median$Age))

#
ggplot(Epithelial_snv_median1, aes(x= Age, y=Ep_vaf, color= Parity)) +
  geom_point(size=0.2) +
  geom_smooth(aes(colour=Parity), method = 'lm', size = 0.2) +
  scale_x_continuous(breaks = round(seq(min(Epithelial_snv_median1$Age),
          max(Epithelial_snv_median1$Age), by = 6),1)) +
  scale_y_continuous(labels = comma_format(accuracy = .01), limits = c(0,1)) +
  scale_color_brewer(palette = "Dark2") +
  labs(x= "Age (yr)", y = "Median VAF \n of SNV")
```

```
## `geom_smooth()` using formula 'y ~ x'
```

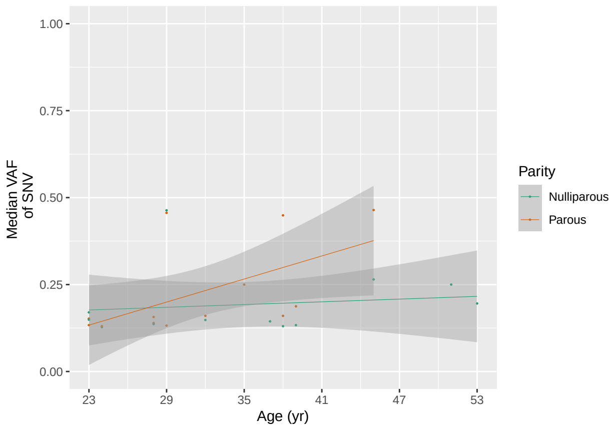


**Plots for indels and complex substitutions were run with the same codes as above, based on the appropriate following subsets.**

```
indels_and_complex<-c("INS", "DEL", "DNP", "TNP" )
Ep_somatic_complex <- Ep_somatic_close[Ep_somatic_close$Variant_Type %in%
          indels_and_complex,]
```

**Violin plots for VAF**

**First define age groups, Early/Late parity (EP/LP), and nulliparity (ENP/LNP) based on 32 as a threshold**

```
Ep_somatic_SNV$Bin <- with(Ep_somatic_SNV, ifelse(
  Age <= 24, '[18-24]', ifelse(
    Age <=28, '[25-28]', ifelse(
      Age <=32, '[29-32]', ifelse(
        Age <=36, '[33-36]', ifelse(
          Age <=40, '[37-40]', ifelse(
            Age <=44, '[41-44]', ifelse(
              Age <=48, '[45-48]', ifelse(
                Age >49, '[>49]', 'NA')))))))))

#plots
ggplot(Ep_somatic_SNV, aes(x= factor(Bin, level = c('[18-24]', '[25-28]',
          '[29-32]', '[33-36]', '[37-40]', '[41-44]', '[45-48]',
          '[>49]')), y=Ep_vaf, fill = Parity)) +
  geom_violin(show.legend = FALSE, size=0.2) +
  stat_summary(fun=median, geom="point", size=0.3, show.legend = FALSE) +
  facet_wrap(~Parity, ncol = 2, scales="free") +
  labs(x= "Age bins", y = "Variant allele fraction \n of SNV") +
  scale_fill_brewer(palette = "Dark2") +
  theme(axis.text.x = element_text(size = 7, angle = 45, hjust=1),
        axis.text.y = element_text(size = 7),
        axis.title.x = element_text(size = 7),
        axis.title.y = element_text(size = 7),
        strip.background = element_blank(),
        strip.text = element_text(size = 7),
        legend.key.height=unit(0.3, "lines"),
        plot.title = element_blank(),
```

```
        panel.grid.major = element_blank(),
        panel.grid.minor = element_blank(),
        panel.border = element_blank(),
        panel.background = element_blank(),
        axis.line = element_line(color = 'black', size = 0.2),
        axis.ticks.length=unit(.05, "cm"))
```

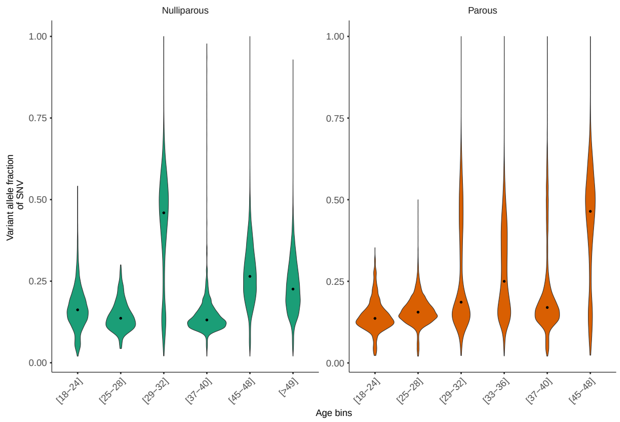


### The same analysis can be done using the same codes on the stromal samples, simply replacing "Epithelial with"Stromal" and "Ep_" with "St_"

#### End
