## Supplementary material for "Mutational landscapes of normal breast during age and pregnancy determine cancer risk": R markdown 2 drivers

B Cereser

### Mutational burden of the normal breast during age and pregnancy

### Analysis of the mutational burden and VAF of breast cancer genes in the epithelium

#### Load Libraries

```
library(data.table)
library(ggplot2)
library(RColorBrewer)
library(scales)
library(dplyr)
```

#### Load in data files

Load in the subset of sample data with only the entry for breast cancer associated genes for the totality of the donors with associated meta-data.
The list of identified breast cancer drivers which we have included in our analysis were taken from the paper "Landscape of somatic mutations in 560 breast cancer whole-genome sequences" by Nik-Zainal et al., Nature, 2016

```
Ep_som_dr <- fread("Ep_som_drivers.txt", sep="\t", header = T)
```

#### Analyse only either NP samples or parous (P) samples where the gap between parturition and tissue collection is $<$ 10 years

This will leave you with the 25 samples analysed in the main paper. The excluded 4 samples are the ones analyzed in the last Extended Data Figure. The column Years_after_pregn (Years after pregnancy) represents the gap between parturition and tissue collection. For Nulliparous (NP) is artificially set at 0 to make it easier when plotting the sets together.

```
Ep_dr_close<-subset (Ep_som_dr, Years_after_pregn <10)
```

### 1. Frequence of samples mutated for each driver gene

#### Calculate how many mutations per gene per sample

```
Ep_dr_close$Ep_Sample_Barcode = factor(Ep_dr_close$Ep_Sample_Barcode)
Ep_dr_close$Hugo_Symbol = factor(Ep_dr_close$Hugo_Symbol)
Ep_dr_close$Parity = factor(Ep_dr_close$Parity)

Ep_dr_mut_count<-Ep_dr_close %>% group_by(Hugo_Symbol,Parity,
         Ep_Sample_Barcode) %>% tally
```

**Calculate how many samples are mutated for each gene, divided by parity**

```
Ep_dr_mut_count$n = factor(Ep_dr_mut_count$n)
Ep_dr_gene_count <- Ep_dr_mut_count[,1:2] %>% group_by(Hugo_Symbol,Parity,
         .drop=FALSE) %>% tally
```

**Divide the subset into the parity groups, to calculate the percentage of mutated samples based on n number of samples.**
N nulliparous = 12, N parous = 13

```
Ep_dr_gene_count_P<-dplyr::filter(Ep_dr_gene_count, grepl('Parous', Parity))
Ep_dr_gene_count_P$Fraction<-Ep_dr_gene_count_P$n/13*100
Ep_dr_gene_count_NP<-dplyr::filter(Ep_dr_gene_count, grepl('Nulliparous', Parity))
Ep_dr_gene_count_NP$Fraction<-Ep_dr_gene_count_NP$n/12*100

Ep_dr_gene_perc<-rbind(Ep_dr_gene_count_P,Ep_dr_gene_count_NP)
Ep_dr_gene_perc$Fraction_facet<-ifelse(Ep_dr_gene_perc$Parity == "Nulliparous",
         Ep_dr_gene_perc$Fraction * -1, Ep_dr_gene_perc$Fraction)
```

**Plots the graph with the frequency of mutated samples**

```
ggplot(Ep_dr_gene_perc, mapping = aes(x = reorder(Hugo_Symbol, Fraction) ,
         y = Fraction_facet)) +
  geom_bar(stat = "identity", position="identity", fill = "grey70") +
  labs(y= "Fraction of \n mutated samples (%)", x = "") +
  coord_flip() +
  geom_hline(yintercept=0, size=0.3)+
  scale_y_continuous(breaks = pretty(Ep_dr_gene_perc$Fraction_facet),
         labels = abs(pretty(Ep_dr_gene_perc$Fraction_facet)), limits = c(-70,70))+
  theme(text = element_text(size=7),
        panel.grid.major = element_line(colour = "Grey95"),
        panel.grid.minor = element_blank(),
        panel.border = element_rect(fill=NA, linetype="solid", colour = "Grey80" ),
        panel.background = element_blank(),
        strip.background = element_blank(),
        strip.text.x = element_text(size = 7),
        axis.line = element_line(color = 'black', size = 0.2),
        axis.ticks.length=unit(.05, "cm"))
```

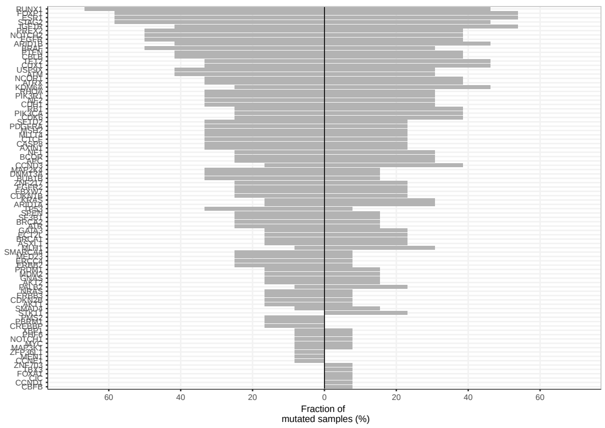

### 2. Heterogeneity of mutations in breast cancer genes

#### Calculate how many mutations per gene per sample

Including the argument .drop=F is useful to determine with samples do not have any mutations in the cancer genes.

```
Ep_dr_mut_count_all<-Ep_dr_close %>% group_by(Ep_Sample_Barcode, Hugo_Symbol,
         .drop = F) %>% tally
```

Two samples, S19 and e37, do not carry any mutations in driver genes, but we want to include them in the graph anyway. **Make a compatible data frame with the missing sample and join it to the previous dataframe and with some extra info on Age and Parity**

```
Ep_Sample_Barcode_ep<-c('S19', 'e37')
n<-c(0,0)
Hugo_Symbol<-c("","")
missing_ep_samples<-data.frame(Ep_Sample_Barcode_ep,Hugo_Symbol,n)
colnames(missing_ep_samples)[1]<-"Ep_Sample_Barcode"
colnames(missing_ep_samples)[3]<-"n"
colnames(missing_ep_samples)[2]<-"Hugo_Symbol"
missing_ep_samples[,3] <- as.numeric(as.character(missing_ep_samples[,3]))

Ep_dr_count1<-bind_rows(Ep_dr_mut_count_all,missing_ep_samples)

sample_age <- fread("clin_info.txt", sep="\t", header = T)
colnames(sample_age)[1]<-"Ep_Sample_Barcode"
sample_age_smaller<-sample_age[,c(1,2:4)]

Ep_dr_count2<-left_join(Ep_dr_count1, sample_age_smaller, by = "Ep_Sample_Barcode")
```

#### Plot stacked bar to represent heterogeneity in the driver gene composition

```
nb.cols <- 95
mycolors <- colorRampPalette(brewer.pal(8, "Set3"))(nb.cols)

#Nulliparous
ggplot(subset(Ep_dr_count2, Parity %in% "Nulliparous"), mapping = aes(
         x = reorder(Ep_Sample_Barcode, Age)  , y = n, fill=Hugo_Symbol)) +
  geom_bar(stat = "identity", colour="black", size=0.07, position = "fill") +
  scale_y_continuous(labels = scales::percent)+
  scale_fill_manual(values = mycolors) +
  labs(y= "mutational burden \n in drivers (%)", x = "Age-ordered nulliparous
         individuals") +
  theme(axis.text.x = element_text(size = 7, angle = 45, hjust = 1),
        axis.text.y = element_text(size = 7),
        axis.title.x = element_text(size = 7),
        axis.title.y = element_text(size = 7),
        plot.title = element_text(size = 7, hjust = 0.5),
        legend.position = "none",
        panel.grid.major = element_blank(),
        panel.grid.minor = element_blank(),
        panel.border = element_blank(),
        panel.background = element_blank(),
```

```
        axis.line = element_line(color = 'black', size = 0.2),
        axis.ticks.length=unit(.05, "cm"))
```

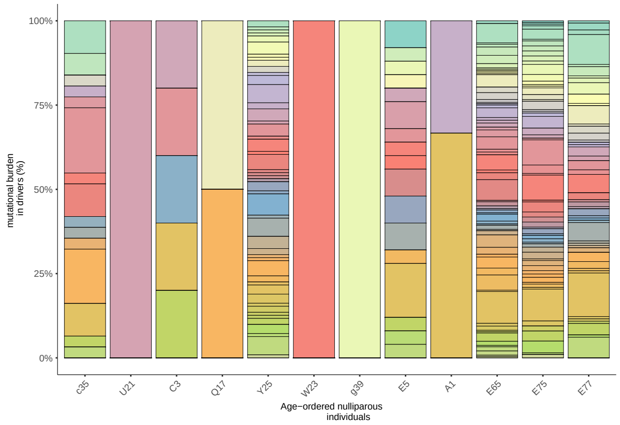


```
#Parous
ggplot(subset(Ep_dr_count2, Parity %in% "Parous"), mapping = aes(
         x = reorder(Ep_Sample_Barcode, Age) , y = n, fill=Hugo_Symbol)) +
  geom_bar(stat = "identity", colour="black", size=0.07, position = "fill") +
  scale_y_continuous(labels = scales::percent)+
  scale_fill_manual(values = mycolors) +
  labs(y= "mutational burden \n in drivers (%)", x = "Age-ordered parous
         individuals") +
  theme(axis.text.x = element_text(size = 7, angle = 45, hjust = 1),
        axis.text.y = element_text(size = 7),
        axis.title.x = element_text(size = 7),
        axis.title.y = element_text(size = 7),
        plot.title = element_text(size = 7, hjust = 0.5),
        legend.position = "none",
        panel.grid.major = element_blank(),
        panel.grid.minor = element_blank(),
        panel.border = element_blank(),
        panel.background = element_blank(),
        axis.line = element_line(color = 'black', size = 0.2),
        axis.ticks.length=unit(.05, "cm"))
```

```
## Warning: Removed 2 rows containing missing values (geom_bar).
```

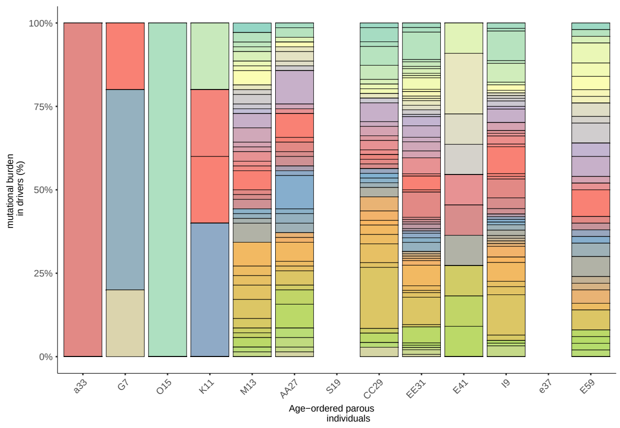


**The same analysis can be done using the same codes on the stromal samples, simply replacing "Epithelial with"Stromal" and "Ep__" with "St__". For the last plot, note that stromal samples which do not contain driver mutations are different and can be defined as:**

```
St_Sample_Barcode_st<-c('D4', 'R18','h40','B2')
```

**End**
